# Robust nuclease-dead S. aureus dCas9-mediated alpha-synuclein knockdown in substantia nigra in a humanized mouse model of Parkinson’s disease

**DOI:** 10.1101/2023.09.05.556425

**Authors:** C. Alejandra Morato Torres, Faria Zafar, Danuta Sastre, De’Angelo Hermesky, Max Y. Chen, Jocelyn Palafox Vazquez, Lei Stanley Qi, Deniz Kirik, Birgitt Schüle

**Affiliations:** Department Pathology, Stanford University School of Medicine, Stanford, CA, USA; SRI International, Menlo Park, CA, USA; Department of Bioengineering, Stanford University, Stanford, CA, USA; Sarafan ChEM-H, Stanford University, Stanford, CA, USA; Chan Zuckerberg Biohub – San Francisco, San Francisco, CA, USA; Department of Experimental Medical Science, Lund University, Lund, Sweden

## Abstract

Parkinson’s disease (PD) is becoming increasingly prevalent due to an aging society, which places a substantial disease burden on patients and their families and an annual cost estimated at 52 billion dollars. However, no approved disease modulatory therapies that halt disease progression are available. Alpha-synuclein is a critical therapeutic target found in aggregated form in Lewy bodies which is the diagnostic hallmark of PD. Familial autosomal dominant forms of PD can present with causative exonic point mutations, copy number multiplications, and non-coding risk variants in the alpha-synuclein gene. The disease onset, severity, and progression depend on the gene expression levels of the alpha-synuclein. Here, we demonstrate that *Streptococcus aureus* dCas9 (sadCas9)-mediated CRISPR interference (CRISPRi) reduces alpha-synuclein mRNA and protein levels in a humanized mouse model. The mechanism of action is based on the principle that a complementary single guide RNA (sgRNA) recruits the sadCas9 protein to the promoter region of alpha-synuclein and modulates target gene transcription, leading to reduced gene expression. We show robust downregulation of alpha-synuclein in neurons after unilateral stereotactic injection into the substantia nigra of adult mice 1 and 6 months after surgery. This work shows proof of concept that viral-mediated sadCas9 CRISPR interference can be a promising therapeutic strategy to reduce alpha-synuclein *in vivo*.

**Graphical abstract:** 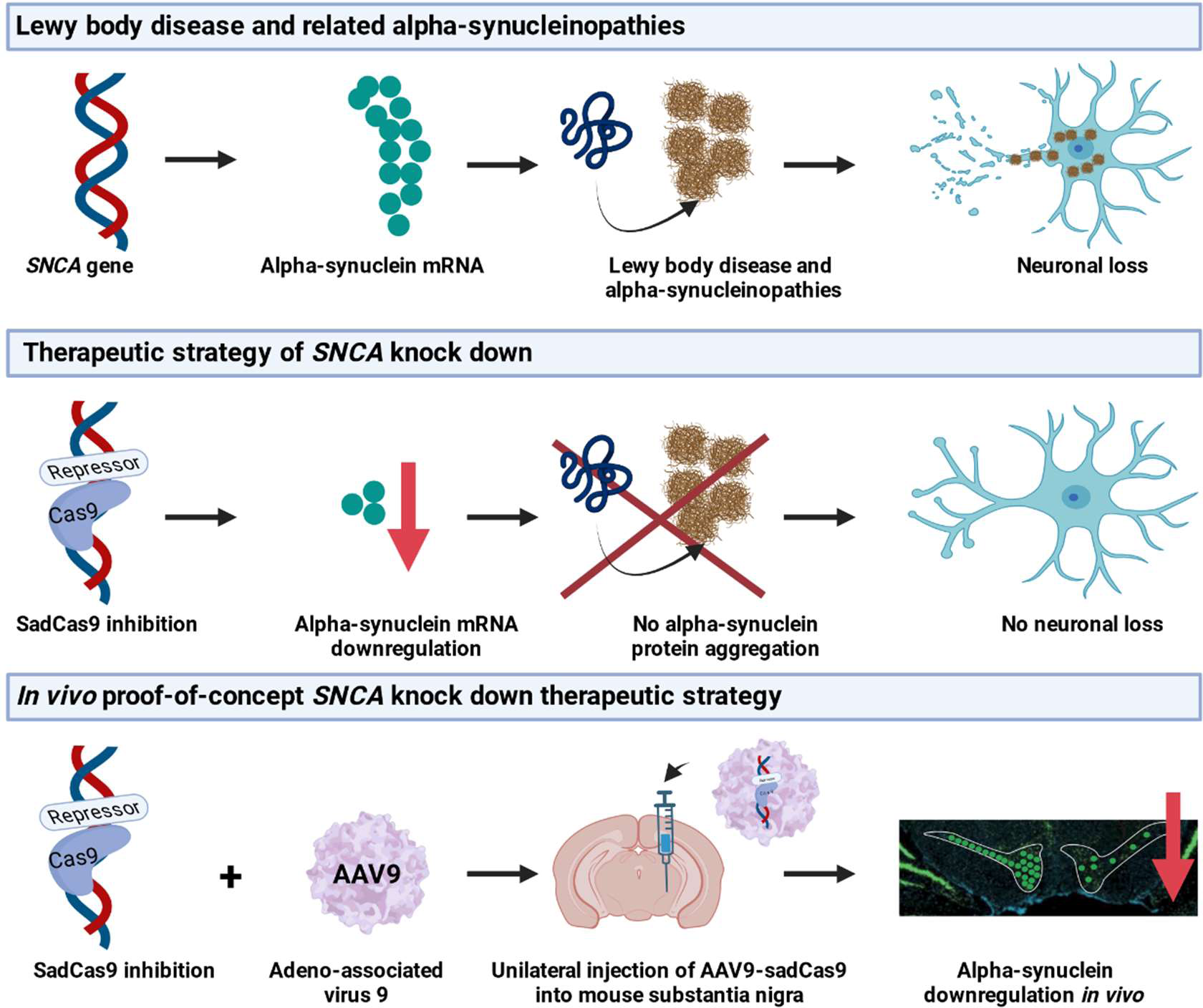

**eTOC synopsis:** This study focuses on the development of an AAV9 nuclease-dead S. aureus CRISPR/Cas9 expression system designed to target the human *SNCA* promoter. The objective is to achieve downregulation of α-synuclein (a-syn) expression. After performing surgery and introducing the expression system, the levels of a-syn were measured at 1 month and 6 months post-surgery. The results indicate a significant downregulation of a-syn at both time points. Furthermore, it was observed that the initial immune response to the system attenuated over time, reaching control levels at the 6-month mark. These findings suggest the potential of this expression system for long-term downregulation of a-syn and provide insights into the immune response dynamics associated with its use.

## Introduction

Parkinson’s disease (PD) is chronic neurodegenerative disease presenting with a varying temporal combination of cognitive, psychiatric, and motor impairments^1,2^. PD is part of the Lewy body disease (LBD) spectrum which encompasses the overlapping diagnoses of PD, Dementia with Lewy bodies (DLB), and PD dementia (PDD). Current treatments are entirely symptomatic, and to date, there are no available therapies proven to cure, halt, or slow disease progression.

Alpha-synuclein is a key therapeutic target and it is found as protein aggregates in intracellular Lewy bodies, which are the diagnostic hallmarks of PD^3^. Furthermore, familial autosomal dominant forms of PD demonstrate pathogenic point mutations, copy number multiplications, and non-coding risk variants in the alpha-synuclein (*SNCA*) gene^4,5^. The disease onset, severity, and progression depend on the gene levels or copy numbers of the *SNCA* gene^6–8^. These studies have led to a widespread hypothesis that lowering alpha-synuclein levels could be of therapeutic benefit for LBD spectrum^9,10^.

Mechanistically, lowering alpha-synuclein levels *in vitro* and *in vivo* has been shown to be neuroprotective and can reverse many PD-related cellular phenotypes.^11–16^ As a proof-of-concept that reduction of alpha-synuclein expression has a neuromodulatory effect, the administration of small interference RNAs (siRNA) in the CNS has been shown to be effective against endogenous murine alpha-synuclein^16–19^. In a translational non-human primate model, siRNA knockdown showed alpha-synuclein reduction in MPTP-exposed squirrel monkeys^13^. A small molecule beta2-adrenoreceptor agonist has been shown to be a regulator of the *SNCA* gene by modifying the histone 3 lysine 27 acetylation of the *SNCA* promoter and improving mitochondrial function in human stem cell-derived neuronal cultures carrying a triplication of the *SNCA* genomic locus^20^. Conceptually, the reduction of alpha-synuclein protein levels has been a widely attempted therapeutic approach and the RNA and protein level with antisense oligonucleotides, small molecules and immune therapy^21^. Several immunotherapeutic approaches are currently being evaluated in clinical trials^22,23^. Small molecules and antisense oligonucleotides that disaggregate toxic alpha-synuclein species or destroy alpha-synuclein mRNA are in late-stage preclinical programs^24^. Lastly, early pre-clinical studies that show proof of concept *in vitro* use the common *S. pyogenes* CRISPR-sadCas9 system for editing or modulation of the *SNCA* gene^25,26^.

New versatile gene-engineering technologies with the power to precisely target positions in the human genome can also be exploited to lower alpha-synuclein mRNA expression level. The Cas9 protein is one of these powerful and now well-known gene engineering technologies that can be used for genomic editing, but also provides features to modulate gene expression by activating or inactivating its expression via steric hindrance of transcription itself or epigenetic changes such as DNA methylation or chromatin modification^27–29^. The first proof that nuclease-dead spdCas9 regulates alpha-synuclein expression was published in 2016^30^. A limited number of single guide (sg)RNAs with a protospacer adjacent motif (PAM) recognition sequence for *S. pyogenes* Cas9 (spCas9) were tested and one sgRNA was found to downregulate alpha-synuclein expression by approximately 50% in cancer cell lines and human induced-pluripotent stem cell (iPSC) models^30^.

In this study, we developed a gene-engineering approach of a smaller nuclease-dead CRISPR/*S. aureus* Cas9 (sadCas9) fused to Krueppel-associated box (KRAB) domains to regulate alpha-synuclein expression and demonstrate that sadCas9 interference technology facilitates alpha-synuclein mRNA and protein reduction in a humanized mouse model with a P1 artificial chromosome (PAC) gene containing the entire human alpha-synuclein gene including introns and their gene regulatory regions^31^. SaCas9 has a gene size of 3.2kb, which is much smaller than that of SpCas9 (4.2kb), which confers a major technical advantage when testing in the challenge-to-delivery cells. We show robust downregulation of alpha-synuclein after unilateral stereotactic injection into the substantia nigra of adult mice 1 and 6 months after surgery. This work shows proof-of-concept that adeno-associated virus (AAV)-mediated CRISPR/sadCas9 interference can be a promising therapeutic strategy to reduce alpha-synuclein *in vivo*.

## Results

### SgRNA in transcriptional start site 2 (TSS2) of human SNCA promoter shows robust downregulation of alpha-synuclein expression in HEK293T cells

Previous studies have shown that targeting of nuclease-dead Cas9 (dCas9) to gene promoter regions can lead to robust and specific gene downregulation due to interference with the gene transcription machinery. This has been achieved by the introduction of inactivating mutations in the Cas9 gene in the two catalytic domains HNH and RuvC which abolishes the nuclease activity, but Cas9 can still be guided to the genomic target sequence, bind to the DNA, and blocks the transcriptional process as an RNA-guided DNA-binding complex^32–35^. When Cas9 is fused to one or more KRAB domains, the downregulation is even more pronounced presumably due to epigenetic changes in histone modification with loss of histone H3-acetylation and an increase in H3 lysine 9 trimethylation in the promoter target region^36,37^.

Based on our previous *in vitro* screen for nuclease-dead *S. aureus* Cas9 (sadCas9) sgRNAs in the human *SNCA* promoter^38^, we tested five sgRNAs around the transcriptional start site 2 (TSS2) of the human *SNCA* gene (TSS2-sg1 to TSS2-sg5) with homology to non-human primates (**Figure 1A,B**). HEK293T cells were triple transfected with the doxycycline-inducible SadCa9-RFP, sgRNA-BFP, and reverse tetracycline-controlled transactivator construct (**Figure 1C**) and yielded a transfection efficiency of 50-60% after FACS sorting (**Figure 1E**). Twenty-four hours post transfection, cells were treated with doxycycline for 48h to activate Cas9 expression (**Figure 1D**) and then further processed with FACS sorting and Taqman analysis. TSS2-sg1, TSS-sg2, and TSSsg4 showed a greater than 50% downregulation compared to the sadCas9 only control (**Figure 1F**).

**Figure 1.**
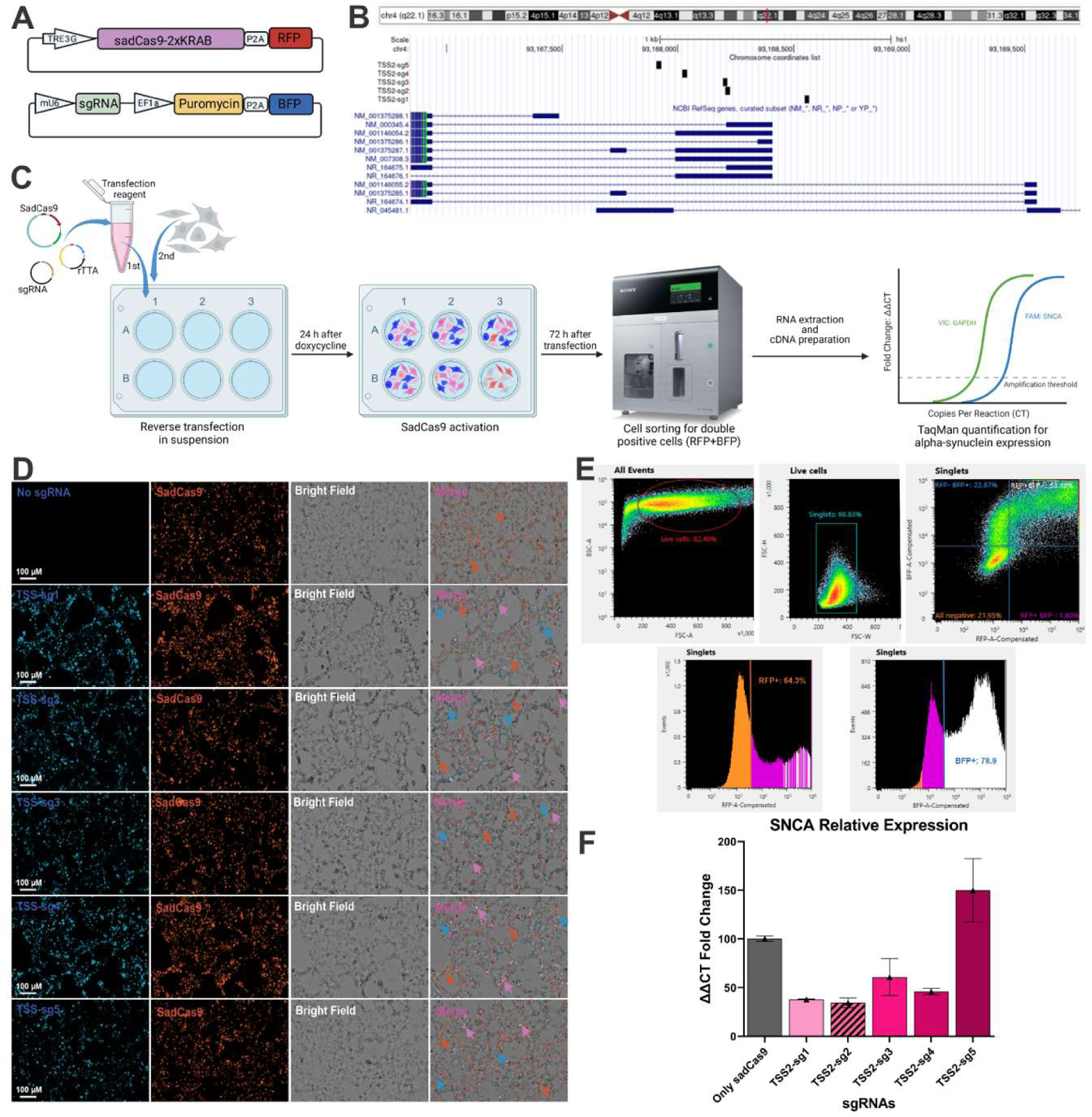
Screening of sgRNAs targeting TSS2 in the *SNCA* promoter with HEK293 cells. (**A**) Schematic representation of vector cassettes for inducible SadCas9 and red fluorescent protein (RFP) and sgRNAs and blue fluorescent protein (BFP) co-expression. (**B**) UCSC Genome browser view of *SNCA* promoter region showing five sgRNAs targeted to the transcriptional start site 2 in the human *SNCA* promoter. (**C**) Schematic overview of the transient inducible expression of sadCas9 and sgRNA, fluorescent activated cell sorting (FACS) and quantitative PCR amplification with Taqman probes. (**D**) Representative images of induced expression of sadCas9 with different sgRNAs and negative SadCas9 only control. (E) FACS sorting and gating of RFP and BFP double-positive cells, on average we reached 53.24% double positive cells in HEK293 cultures. (F) Taqman quantitative expression analysis of the *SNCA* gene, normalized to GAPDH, and compared to sadCas9 only cells.

For *in vivo* experimentation, we choose sgRNA TSS2-sg2 (chr4:93,168,206-93,168,226 (T2T CHM13v2.0/hs1)) because of its strong downregulation and 100% homology to the non-human primate *SNCA* promoter region (rhesus macaque, squirrel monkey, and cynomolgus), and no predicted off-targets for up to 4 mismatches in the human genome reducing the chance off-target effects.

### AAV cassette design for co-expression of sadCas9 and TSS2-sgRNA2 in vivo

The challenge with AAV genomes is their small packaging capacity of approximately 4.7 kb, including the ITR repeats^39^. To not exceed the AAV packaging capacity, but to include all elements into one vector, we chose the sadCas9 variant. SadCas9 has a length of 3.2 kb and is about 1 kb shorter than the more commonly used spCas9. We selected the MECP2 promoter, which is small in size (229 bp)^40,41^ compared to synthetic promoters such as cytomegalovirus (CMV) or CAG (CMV enhancer fused to chicken beta-actin promoter) promoters and has been used in gene therapy strategies^42^. Next, we placed the human U6 promoter which is 249bp in length and is 65 bp shorter than the mouse U6 promoter upstream of the sadCas9 constructs^43^. A guanosine nucleotide follows the human U6 promoter to initiate transcription of the TSS2 sgRNA 2^44^ and sadCas9-optimized scaffold for binding to the *SNCA* promoter target sequence. With this design the expression cassette has a size of 4,796 bp.

We chose an AAV9 serotype, which has been well-investigated for delivering transgenes in the CNS, is a compelling candidate for gene therapy in humans. The AAV9 delivery vector used for our studies has been used in several Parkinson’s disease clinical trials and shows a favorable safety profile^45,46^

### *In vivo* study design for human alpha-synuclein downregulation in PAC animal model

The *in vivo* study design consisted of six groups of four months old adult mice at time of surgery (**Figure 2**). Two groups received 1 µl of TSS2-sgRNA2 AAV at a titer of 1.50 x 10 ^10^ genome copies/µl stereotactically injected into the substantia nigra unilaterally and mice were euthanized one or six months after surgery, two groups received 1 µl of control sgRNA which is a non-targeting sgRNA, and two groups received 1 µl of saline solution as controls (**Figure 2**).

**Figure 2.**
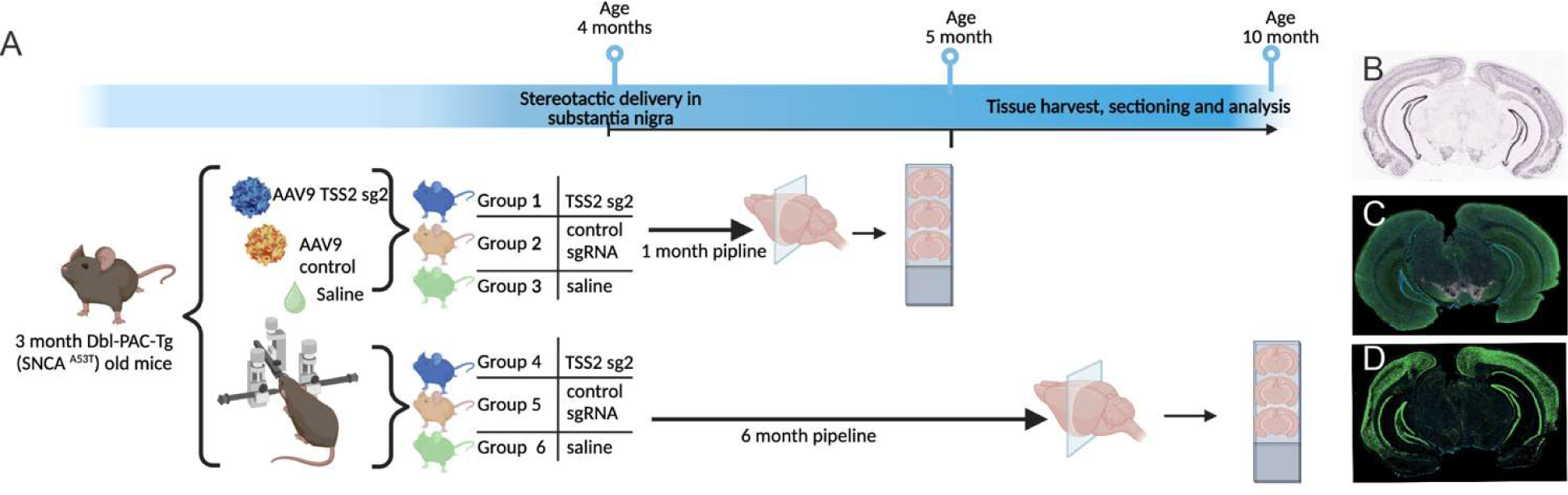
Study design and timeline of experiments. (**A**) Four months old Dbl-PAC-Tg (SNCA^A53T^) mice were grouped into six experimental groups, groups 1 and 4 received AAV9 virus containing the active construct of sadCas9 with TSS2 sgRNA2 stereotactically injected into the substantia nigra groups 2 and 5 received AAV9 containing a non-human targeting sgRNA gal4 control, and groups 3 and 6 received saline controls. Groups 1-3 were euthanized after 1 month and groups 4-6 after 6 months and brains and peripheral organs were harvested and fresh frozen in liquid nitrogen. Created with Biorender.com. (**B**) Coronal sections of 20 um were taken and further stained by in situ hybridization RNAScope or (**C**) immunofluorescence (alpha synuclein: green and Tyrosine Hydroxylase: white) histology, quantified, and analyzed by Qpath (v 0.4.2). (**D**) Comparative in situ hybridization mRNA for mouse alpha synuclein from Allen Mouse Brain Atlas; experiment-RP-Baylor_103043-coronal. Created with Biorender.com

No adverse effects of the treatment were observed in any of the cohorts. Throughout the study, the overall health and development of the animals in the treatment groups remained consistent with that of sham-treated mice. The mice gained weight over time (**Supplemental Figure 1**) with no difference between the experimental groups, and no evident behavioral abnormalities were noted. Following euthanasia, the gross macroscopic appearance of all internal organs during necropsy was normal in all mice.

### Unilateral stereotactic delivery of AAV9 virus vector shows widespread expression of sadCas9

We detected sadCas9-positive staining spreading on the ipsilateral site of injection in the substantia nigra to the cortex, hippocampal region, substantia nigra, thalamus and hypothalamus (lateral (LMN) and medial mammillary nucleus (MMN)). Cas9 expression was localized diffuse in the cytoplasm and nucleus with prominent expression in the substantia nigra pars compacta and pars reticulata (SNpc, SNpr), ventral tegmental area (VTA) and lower expression in fiber tracts towards the lower cerebral peduncle. Moreover, Cas9 expression colocalizes with the tyrosine hydroxylase (TH), a key marker for dopaminergic neurons (**Supplemental Figure 2**). At 6 months post-injection, there is a decrease in sadCas9 signal for mRNA and protein expression. However, we did not detect neuronal cell loss at 1 or 6 months post stereotactic injection (**Supplemental Figure 3**).

### Robust downregulation of human alpha-synuclein mRNA and protein in the substantia nigra

The main objective of this study was to provide *in vivo* evidence showcasing the feasibility of our previously established *in vitro* strategy for downregulating alpha-synuclein. To detect SNCA mRNA, we employed an in situ hybridization probe specifically designed for human SNCA mRNA (RNAScope). We confirmed that the spatial distribution pattern of human SNCA mRNA in our PAC transgenic mouse model aligned with the expression pattern of the mouse Snca mRNA documented in the Allen Brain Atlas (http://mouse.brain-map.org/) (**Figure 2**).

We analyzed mRNA expression in coronal sections of the mesencephalon, both at one and six months following stereotactic surgery. Our analysis revealed a reduction in RNA foci in the TSS2 sgRNA2 group compared to the two control groups. Specifically, one month after the surgery, a substantial reduction in SNCA mRNA levels was observed, averaging a 73% decrease when compared to the saline control group. (**Figure 3A, B**). These findings illustrate that sadCas9 guided by TSS2-sgRNA2 led to efficient suppression of SNCA mRNA expression in the substantia nigra. (**Figure 4A**). The effect of SNCA mRNA downregulation persisted at the 6-month post-injection, the ipsilateral site displayed a significant average decrease of 63% in SNCA mRNA levels. (**Figure 4B**). There was no statistically significant difference in the mRNA downregulation of TSS2-sgRNA2 between the both time points, as indicated by the RNAScope signal.

**Figure 3.**
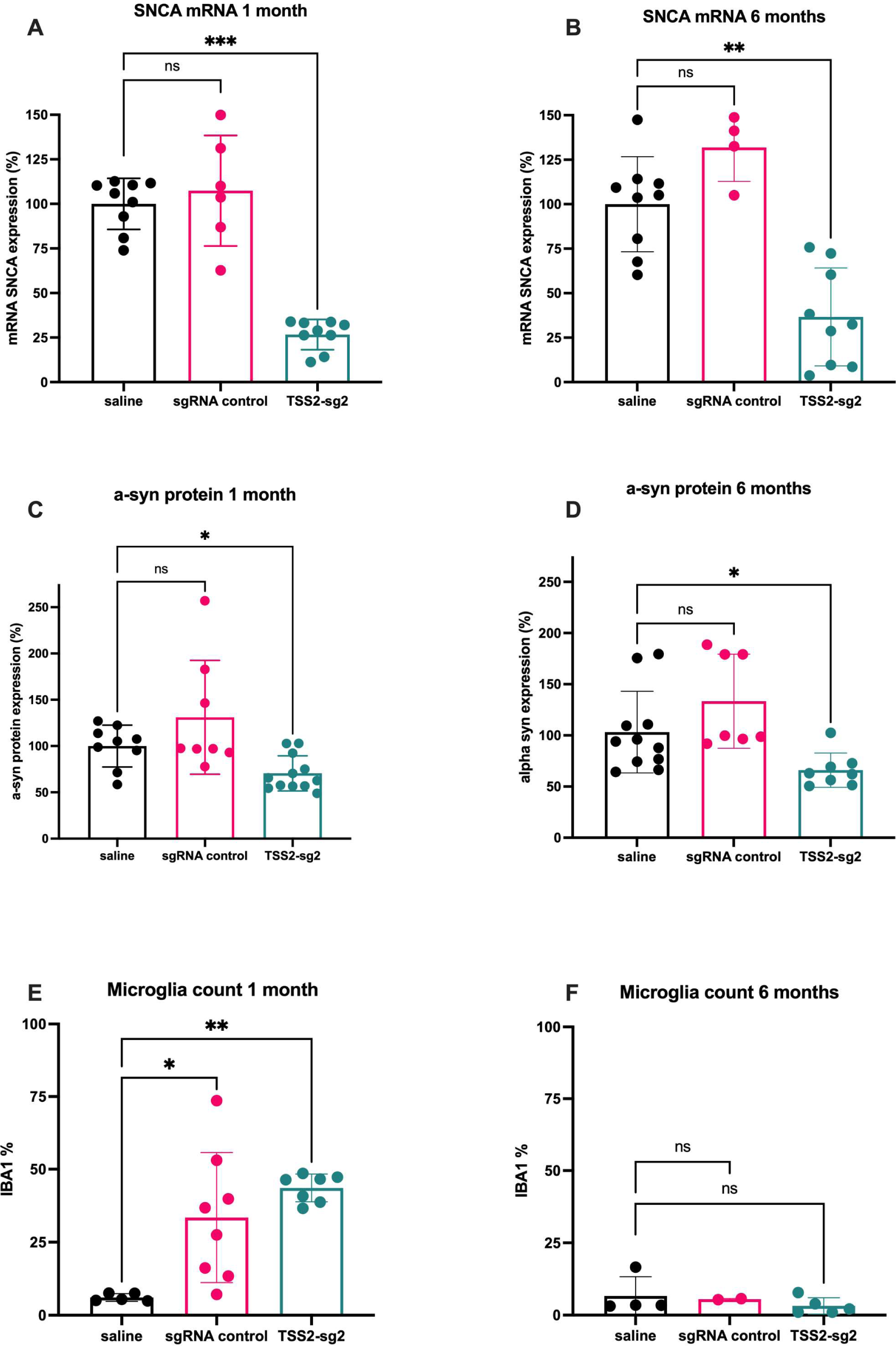
Graphic representation of alpha synuclein mRNA, protein, and immune response after AAV9-mediated SadCas9 interference. **(A)** Graphic representation of *SNCA* mRNA expression at 1 month post injection and **(B)** 6 months post-injection, **(C)** graphic representation of alpha synuclein protein at 1 month and **(D)** 6 months post injection. **(E)** immune response measured by microglial population at 1 month and **(F)** 6 months post-injection. Results are shown as ns = non-significant, (*) p<0.05, (**) p< 0.01, or (***) p< 0.001.

**Figure 4.**
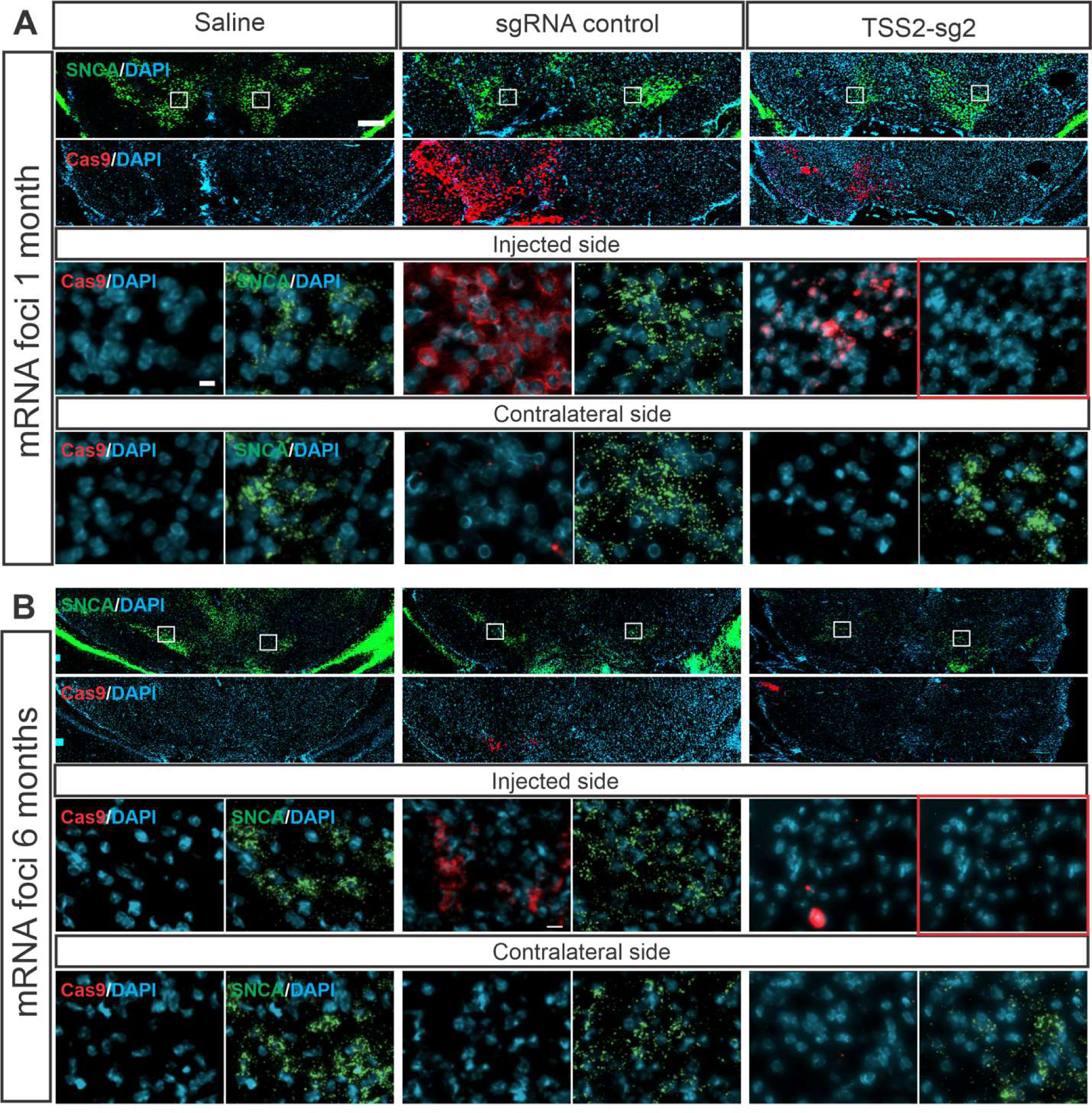
Robust *SNCA* mRNA downregulation 1 and 6 months after AAV9-mediated SadCas9 interference. **(A)** *SNCA* mRNA downregulation at 1-month post-injection and **(B)** *SNCA* mRNA downregulation at 6 months post-injection. RNA expression is detected using specific hybridization probes (RNAScope) against alpha-synuclein (green) and cas9 (red) counterstained with DAPI. Left panel: Saline control, middle panel; sgRNA control and right panel TSS2-sg2 group. Injected side and contralateral side are represented for comparison. Red squares in TSS2-sg2 indicate the mRNA alpha synuclein reduction at 1 and 6 months post-injection. Scale bars at 200 µm and 10 µm.

Next, we evaluated alpha-synuclein downregulation at the protein level using immunofluorescence in the same region with coronal sections of the mesencephalon. Immunofluorescence signal for alpha-synuclein protein showed a downregulation of 37% at 1 month and 32% at 6 months when compared to the saline control group (**Figure 3C, D; Figure 5**). There was no statistically significant difference in the alpha-synuclein protein downregulation of TSS2-sgRNA2 between the both time points, as indicated by the immunofluorescence signal.

**Figure 5.**
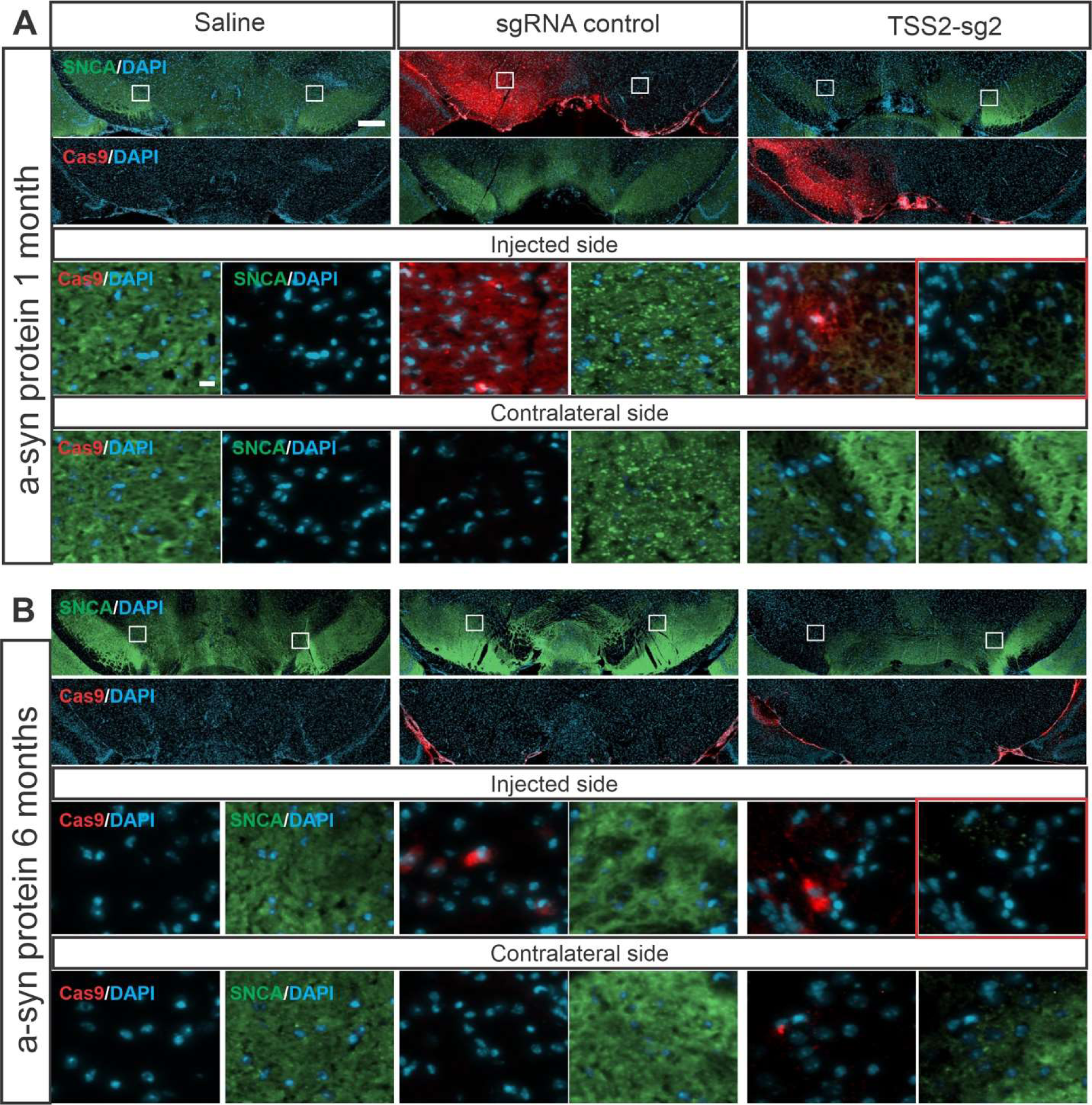
SadCas9-mediated alpha-synuclein protein downregulation at 1 and 6 months post stereotactic surgery in substantia nigra. **(A)** Alpha synuclein protein downregulation at 1-month post-injection and **(B)** Alpha synuclein protein downregulation at 6 months post-injection. Protein detection is performed by immunofluorescence with a human alpha synuclein (green) specific antibody and a Cas9 (red) antibody and counterstained with DAPI. Left panel: Saline control, middle panel; sgRNA control and right panel TSS2-sg2 group. Injected side and contralateral side are represented for comparison. Red squares in TSS2-sg2 indicate the alpha synuclein protein reduction 1 and 6 months post-injection. Scale bars at 200 µm and 10 µm.

The observed discrepancy between mRNA and protein levels of alpha-synuclein could potentially be elucidated by considering the inherent characteristics and subcellular distribution of the alpha-synuclein protein in contrast to its corresponding RNA. The SNCA mRNA is predominantly detected in the perinuclear region, manifesting as discrete punctate structures. Conversely, the alpha-synuclein protein exhibits a more widespread presence throughout the entire cell and is particularly enriched in presynaptic regions. Notably, the protein detection was conducted within the SNpc and VTA regions, encompassing the capture of alpha-synuclein signals originating from diverse cell types or projections originating from other cerebral areas. This might contribute to the dilution effect of the alpha-synuclein signal.

### Attenuation of microglia immune response six months after surgery to control levels

Despite the CNS’s limited clearance of AAV (adeno-associated virus) by immune responses, the issue of neuroinflammation is progressively assuming significance within the domain of AAV-mediated gene delivery. The administration of AAV specifically to the CNS has been correlated with a notable upsurge in neutralizing antibodies, along with the circulation of diverse immune cell subtypes, encompassing CD45+ leukocytes, natural killer cells, as well as CD8+ and CD4+ T lymphocytes.

As part of our study, we focused on examining potential upregulation of IBA1 expression. IBA1, a constituent of the calcium-binding protein cluster, is a key protein marker to signify microglial activation and also known as allograft inflammation factor 1 (AIF-1), microglia response factor (MRF-1), or daintain. IBA1 plays a pivotal role in orchestrating the restructuring of the microglial cytoskeleton, thereby bolstering the phagocytic processes^47,48^.

One month post stereotactic surgery, besides a substantial upregulation of Cas9 protein expression at the site of injection, we also detected a pronounced augmentation in the number of IBA1-positive cells, demonstrating an average increase of 27%. This pattern was pronounced in both the TSS2-sg2 RNA and control sgRNA groups, relative to the saline control group (**Figure 3E**, **Figure 6A**). The increased number of IBA1-positive cells serves as a strong indicator of microglia activation at the injection site compared to both the contralateral side and the control animals that received saline injections. This observation suggests that the expression of Cas9 protein causes an activation of microglia.

**Figure 6.**
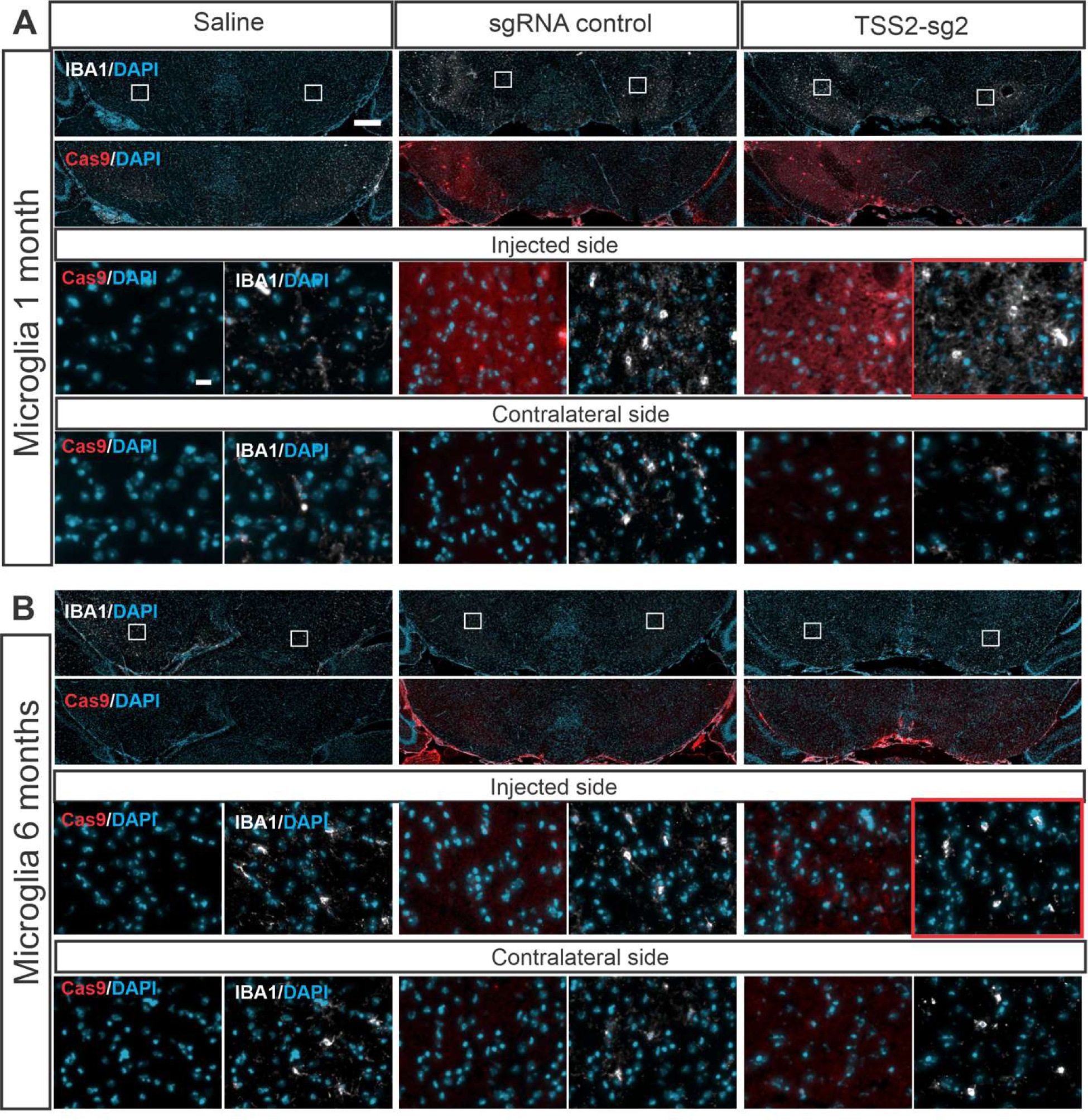
Immune response attenuated post-stereotactic surgery in substantia nigra. **(A)** Immune response at 1-month post-injection and (**B)** Immune response at 6-month post-injection Immune response was analyzed by percentage of microglia in the substantia nigra and ventral tegmental area at 1- and 6-months post-injection. Microglia marker: Iba-1 (white) and Cas9 (red) were revealed by immunofluorescence and counterstained with DAPI. Left panel: Saline control, middle panel; sgRNA control and right panel TSS2-sg2 group. Injected side and contralateral side are represented for comparison. Red squares in TSS2-sg2 1 month indicate the increased microglial response and at 6 months the microglia population reversed to baseline. Scale bars at 200 µm and 10 µm.

However, this increase of IBA1 signal was transient and returned to saline control levels by the sixth month post-injection. This suggests that the microglial activation triggered by the injection is temporary and returns to baseline (**Figure 3F**, **Figure 6B**).

## Discussion

Alpha-synuclein stands as a prominent target for novel therapeutic approaches in addressing Parkinson’s disease (PD) and related synucleinopathies. The significance of targeting alpha-synuclein stems from evidence that increased expression of the wildtype protein alone can trigger neurodegeneration, evident in patients with SNCA genomic locus duplications, triplications, and common non-coding alterations that can elevate its expression. This heightened expression’s link to neurodegeneration and disease severity supports its causal role. Clinical genetic findings and *in vitro* and *in vivo* studies have spurred the notion that diminishing alpha-synuclein content or eradicating toxic forms could potentially mitigate, reverse, or even halt disease progression. Various strategies have emerged, including lowering mRNA via RNA interference, hindering aggregation, fostering intracellular degradation, enhancing extracellular breakdown through immunotherapies, and reducing uptake of extracellular alpha-synuclein. Prominently, alpha-synuclein antibodies have progressed to human clinical trials, with their potential to ameliorate PD symptoms currently under assessment.

This study is built on two previous publications that showed evidence for proof of concept that the *SNCA* promoter can be successfully targeted by CRISPR/Cas9 interference^30,49^. Heman-Ackah et al. found one sgRNA in the region of the transcriptional start site 2 (TSS2) of the *SNCA* promoter that showed a 50% *SNCA* downregulation in HEK293 cultures and human iPSC-derived neurons. In that study, the *S. pyogenes* version of CRISPR/Cas9 was used which is 4.2 kb in length. In our previous *in vitro* study^49^ and in this study, we used *S. aureus* dCas9 guided to the promoter region of *SNCA* gene which also robustly mediated reduction of mRNA and protein levels *in vitro*. SadCas9 is one of the smaller Cas9 proteins which are developed for *in vivo* gene therapy applications. We also showed that lowering alpha-synuclein in human iPSC-derived neuronal cultures from a patient with an SNCA triplication is reducing mitochondrial DNA damage and oxidative stress levels suggesting a reduction in disease burden^38^.

### In vivo gene editing approaches using CRISPR/Cas9 in AAV virus and its immune response

*In vivo* demonstrations of CRISPR technology’s proof-of-concept span heart disease, HIV, and other conditions^50–52^. However, a recent N-of-1 human clinical trial, a patient received a systemic delivery of rAAV9 *CRISPR/Cas9*-transactivator for advanced Duchenne muscular dystrophy and died six days after treatment of severe acute-respiratory distress syndrome with diffuse alveolar damage due to systemic innate immune toxicity^53^. This case illustrates the concern that AAV-mediated gene therapy heightens the potential for critical innate and adaptive immune responses directed at the AAV vector capsid and the transgene product^54^. Studies have demonstrated that AAV therapies may induce immune responses that vary depending on the dose administered, ultimately impacting the therapeutic effectiveness if not properly identified and effectively managed^55^. Consequently, it is crucial to address these immune responses optimally to ensure the success and safety of AAV-based gene therapies.

In our method, we opt for targeted administration of the AAV9 system specifically into the substantia nigra, a key region for motor symptoms in Lewy body disease. This approach allows us to observe virus spread both in retrograde and anterograde directions. Unlike systemic delivery, we introduce a limited virus quantity, aiming to reduce the immediate immune reaction. Despite detecting an activated immune response characterized by heightened microglia activity, we note that this response subsides within six months without causing neuronal cell loss or other adverse effects.

Viral vectors are designed to deliver a DNA payload efficiently; however, the human immune system’s innate pathogen-associated molecular pattern (PAMP) sensors can trigger adaptive effector functions. Unmethylated CpG dinucleotide-based motifs are known PAMPs that bind and dimerize monomeric toll-like receptor 9 (TLR9) expressed in human dendritic cells and activate signaling pathways, thereby promoting cytotoxic T-cell (CTL) responses to AAV vectors^56^. To further optimize the sadCas9 expression cassette for clinical translation in future pre-clinical studies, we aim to reduce this CTL response by codon-modification of the sadCas9 in the AAV9 expression cassette to remove CpGs.

### How much Cas9 expression is necessary for long-term downregulation of alpha-synuclein?

After AAV delivery, we observed robust SadCas9 expression one month post-surgery. However, this expression dwindled by the six-month mark, despite the ongoing downregulation of alpha-synuclein. Theoretically, the targeting of both alleles of the SNCA gene per cell requires only a pair of SadCas9 molecules guided by specific sgRNAs. And excessive SadCas9 expression might have counterproductive effects, as it could sequester factors of the transcriptional machinery, binding to free SadCas9 molecules.

Another mechanism of sustained downregulation could be an epigenetic effect of the Krueppel-domain modifying histone marks or mitigating CpG methylation. A similar Cas9 strategy with a fused with the catalytic domain of DNA-methyltransferase 3A (DNMT3A) to hypermethylate intron 1 shows a 30% downregulation of alpha-synuclein and rescue of mitochondrial reactive oxygen species production and cellular viability *in vitro*^25^.

If achieving sustained alpha-synuclein downregulation requires less SadCas9 than initially assumed, there’s potential for reducing the AAV9 dosage. This reduction in AAV9 quantity could subsequently mitigate immunological side effects and enhance overall safety of the treatment. This will be tested future pre-clinical efficacy studies.

### Regulation of alpha-synuclein to physiological ‘protective’ levels

In comparison to other alpha-synuclein lowering strategies, we propose to target alpha-synuclein at the genomic level. Other therapeutic strategies target either alpha-synuclein mRNA or protein. With our approach, alpha-synuclein is prevented from being transcribed and translated, hence less protein is produced. A crucial facet of our CRISPR/Cas9 SNCA interference approach is its capacity to modulate the alpha synuclein transcript levels by selecting different sgRNAs, allowing for adjustments of protein expression towards specific physiological ranges, e.g. a patient with an SNCA triplication might need a stronger sgRNA than a patient with an SNCA promoter variant.

A critical question is how much alpha-synuclein downregulation will be beneficial. While alpha-synuclein knockout mice are viable and fertile^57^, individuals with SNCA genomic deletions present with neurodevelopmental delay and autism spectrum disorder^4,5^. Our therapeutic strategy is to deliver sadCas9 together with a small guide RNA via gene therapy. It is based on a single intervention principle. Furthermore, this gene targeted approach addresses a direct cause of PD as opposed to all currently available forms of symptomatic treatment. Lowering alpha-synuclein to therapeutically safe levels should be neuroprotective in human which we have been testing in human iPSC-derived neuronal cultures.^11,12,20^

In summary, this study establishes the viability of viral-mediated SadCas9 interference as a promising therapeutic avenue for *in vivo* alpha-synuclein reduction. The persistent decrease in SNCA mRNA expression underscores the success of our *in vivo* intervention, suggesting a possible enduring influence on mitigating alpha-synuclein-associated pathophysiological mechanisms. We hold the perspective that this approach has the potential to modify disease progression and markedly enhance symptom treatment beyond existing standards of care.

## Materials and methods

### Cloning of CRISPR/sgRNA lentiviral constructs with fluorescent selection markers

The *S. aureus* dCas9 in a lentiviral vector is driven by a tetracycline-inducible promoter (TRE3G). To facilitate selection of cells by FACS, pHR:TRE3G-SadCas9-2xKRAB-p2a-RFP was subcloned from a pHR:TRE3G-SadCas9-2xKRAB-p2a-zeo (A gift from Professor Stanley Qi), where zeocin resistance gene (zeo) was replaced by a red fluorescent marker, turboRFP. Briefly, backbone was digested with BamHI and NotI to linearize vector. SadCas9-2xKRAB-p2a segment was amplified by PCR from backbone using high-fidelity polymerase (Phusion, NEB, Cat. No. M0530S); and TurboRFP was PCR amplified using high-fidelity polymerase. Assembly PCR primers were designed to include at least 40 bp of homology with adjacent fragments. Digested backbone and PCR amplified fragments were joined via seamless cloning (NEBuilder® HiFi DNA Assembly Master Mix, NEB, Cat. No. E2621S). Assembled plasmid was Sanger-sequenced to ensure correct assembly and reading frame. The reverse tetracycline-controlled transactivator was expressed from pCMV-rtTA (rtTA-N144), a gift from Andrew Yoo (Addgene # 66810).

#### HEK cell culture

HEK293 cells were maintained in DMEM High Glucose (Cytiva, Cat. No. SH30022.02) supplemented with 10% FBS (Phoenix Scientific, Cat. No. PS-100-02-500), 1X MEM Non-Essential Amino Acids (NEAA) Solution (Thermo Fisher, Cat. No. 11140050), 1X Sodium Pyruvate (Thermo Fisher, Cat. No. 11360070), 1X Glutamax (Thermo Fisher, Cat. No. 35050061) and 50 U/mL Penicillin-Streptomycin (Thermo Fisher, Cat. No. 15070063). Cells were passaged by enzymatic treatment with TrypLE (Thermo Fisher, Cat. No. 12604021) when 90-100% confluent. 6-well plates were coated with 0.1% gelatin for at least 15 mins at room temperature prior to seeding cells. Cells were subcultured at 1.25 - 5 x10^5^ cells/well or split at 1:10 ratio in a 6-well plate.

### Transient reverse transfection

At 80-90% confluence, HEK293 cells were harvested the day of transfection. One 6-well plate can yield 3 - 5 x10^6^ cells. A maximum of 4.5 μg of plasmid (1.5 μg each): pTRE-sadCas9-2xKRAB-2A-RFP (sadCas9-RFP), pCMV-rtTA, and pHR: mU6-sgRNA-EF1A-puro-P2A-BFP (sgRNA-BFP) (1:1:1) using 24 μL TransIT®-LT1 Transfection Reagent (Mirus Bio, Cat. No. MIR 2300) in a final volume of 1000uL Opti-MEM medium (ThermoFisher, Cat. No. 31985062) were combined and incubated for 20 mins, then added to one well of a 6-well plate. Five different sgRNAs (TSS2-sg1, TSS2-sg2, TSS2-sg3, TSS2-sg4, and TSS2-sg5) and one negative control without any sgRNA were used to screen for alpha-synuclein downregulation (**Figure 1, sgRNA table**). Each sgRNAs were used to reverse transfect in each well of a 6-well plate. After adding, the plasmid mixes in a well, 1.25 x10^5^ cells/cm^2^ were immediately added to each well with HEK293 maintenance media modified with 10% tetracycline-free FBS (Takara Bio, Cat. No. 631367). After 24 h, medium was changed to HEK293 media (with Tet-free FBS) supplemented with 500 ng/mL doxycycline to activate SadCas9 expression.

### Fluorescence activated cell sorting (FACS)

Both RFP and BFP positive HEK293 cells were imaged (automated imaging system, ImageExpress Pico) and collected after FACS for RNA extraction 72 h post-transfection/48 h after addition of doxycycline. Briefly, on the day of sorting, transfected HEK293 cells were dissociated using enzymatic treatment with TrypLE for 5 min, centrifuged at 1500 rpm for 5 min, and re-suspended in 1-2 mL of cold PBS. Cell suspensions were filtered through a 100 µM nylon cell strainer (Fisher Sci, Cat. No. 0877123) before acquisition on a flow cytometer (Sony, Model SH800). Sorter was equipped with three excitation lasers: 488 nm, 405 nm, and 561 nm; sorter channels used were FL1 450/50 (BFP), FL2 585/30 (FITC), and FL3 617/30 (RFP). All cells were identified at a sensor gain of FSC 3 and BSC 24%, single cells were gated with FSC-W and FSC-H, and filter voltages were set at 38% FL1, 29% FL2, and 38.5% FL3 based on positive fluorescence signal.

Cells were sorted based on double positive expression of RFP (a marker for SadCas9 expression) and BFP (a marker of sgRNA). Using the 100 μm sorting chip at 30.5 kHz, an average threshold of 10,000 events per second, cells were sorted for FL1/BFP and FL3/RFP (FL2/FITC was used for autofluorescence compensation to allow background signal subtraction) with purity setup. Sorted cells were collected in PBS, centrifuged at 5000 rpm for 5 mins at 4C, and stored pellet for RNA extraction at −080C.

### RNA extraction, cDNA synthesis, and qPCR

RNA extraction and DNase treatment were performed using the PureLink™ RNA Mini Kit (Thermo Fisher, Cat. No. 12183025) following manufacturer’s guidelines. RNA concentration and quality were determined using the NanoDrop Technologies ND-1000 spectrophotometer (Thermo Fisher). cDNA synthesis was completed using 1 ug of RNA as input for random primer reaction (High Capacity cDNA Reverse Transcription Kit; ThermoFisher, Cat. No. 4368814) following manufacturer’s guidelines. cDNA was diluted with nuclease-free water to a concentration of 10 ng/µL. For qPCR, 10 ng of cDNA was used as template. TaqMan Fast Advanced Master Mix (Thermo Fisher, Cat. No. 4444556) and TaqMan probes for FAM-*SNCA* (Thermo Fisher, Assay ID: Hs00240906_m1) and VIC-*GAPDH* (Thermo Fisher, Assay ID: Hs99999905_m1) were used to amplify cDNA. qPCR reactions were run in a QuantStudio 6 Flex Real-Time PCR system (Thermo Fisher, Cat. No. 4485691) and Ct values were obtained using built-in software. NTC samples were included in each plate. RT-control samples were included for each round of RNA extraction and consisted of a cDNA reaction without addition of reverse transcriptase. NTC and RT controls did not amplify in any of plates included in the data presented. Relative mRNA expression was calculated using 2^−ΔΔCt^ method ^58^ using *GAPDH* as a reference gene and Excel software for calculations. The calibrator sample was sgRNA-Gal4, a sgRNA that does not target any human gene. All samples were run in three technical qPCR replicates. A minimum of two biological replicates were used for each sgRNA tested.

### AAV virus production

Plasmid AAV9 was synthesized and cloned, and virus production was performed under a service contract with VectorBuilder, Inc., Chicago, USA. For AAV production and QC control, the transfer plasmid carrying sadCas9/hU6 guideRNA expression cassette was co-transfected with a Rep-cap plasmid and helper plasmid encoding adenovirus genes (E4, E2A and VA) that mediate AAV replication in HEK293T cells. After a short incubation period, viral particles were harvested from supernatant and concentrated by polyethylene glycol (PEG) precipitation. For the in vivo grade ultra-purified AAV, viral particles were further purified and concentrated by cesium chloride (CsCl) gradient ultracentrifugation at >10^13 GC/ml in PBS buffer. A qPCR-based approach measured AAV titer. The quality control includes titer measurement, sterility testing for bacteria and fungi, and mycoplasma detection. For storage and handling, the AAV was stored in PBS-based buffer at −80°C for long term storage.

For stereotactic injections, we thawed a vial of AAV on ice prior to use, prepared a 1.50 x 10 ^10^vg/µl dilution with PBS and kept it on ice during the experiment. We kept thawed and diluted AAV at 4°C for 1 to 2 days when additional surgeries were planned.

### Rodent model for in vivo studies: human transgenic alpha-synuclein mouse model (Dbl-PAC-Tg(SNCAA53T); Snca−/−)

All experiments involving mice were conducted according to animal welfare regulations under protocol Stanford protocol APLAC-33261. Animals were maintained in a controlled facility in a 12-hour dark/12-hour light cycle, a standardized room temperature (RT) of 21 °C, with free access to food and water.

For in vivo experiments, we used a previously established humanized mouse model for alpha-synuclein Dbl-PAC-Tg(SNCA^A53T^);Snca−/− on a mixed genetic background 129S6/SvEvTac * FVB/N and mouse alpha-synuclein knockout ^31^ (http://www.informatics.jax.org/allele/MGI:4412066). The model expresses the human alpha-synuclein gene solely under the genomic regulatory elements of the human *SNCA* promoter (34kb upstream of the *SNCA* gene and all intronic regions that allows splicing). The PAC-Tg(SNCA^A53T^) transgene located on the 146 kb RPCI-1 human male P1 artificial chromosome (PAC) clone 27M07 and contains the entire human *SNCA* (synuclein, alpha (non A4 component of amyloid precursor)) gene and 34 kb of its upstream region and have the p.A53T human mutation. The targeting vector for the *Snca* knockout allele was designed to replace exons 4 and 5 with a reverse-oriented neomycin resistance cassette. The construct was electroporated into 129S6/SvEvTac-derived TC-1 embryonic stem cells. The transgenic dbl-PAC-Tg(SNCA^A53T^) strains and the Snca−/− strain were combined to form a homozygous double transgenic.

### Stereotactic surgery

We anesthetized mice in an induction chamber (SOMNO-0705) with 3-4% isoflurane and used a nose cone to sustain isoflurane anesthesia at 1 - 2% throughout the surgery. When fully anesthetized, we placed the animal in a stereotaxic frame (KOPF, 00512R) on a temperature-controlled heating pad (S) and applied ophthalmic ointment (CLC medica, 12311) to keep eyes moist during surgery and prevent cornea damage. We then shaved head and neck of the mouse with a razor. Exposed skin was cleaned with a 70% alcohol wipe and then two times with iodine solution (2%). Skull was exposed using a #10 scalpel (Integra, 12460451E) and a single hole was drill in the AP, ML coordinates. Coordinates used for SN injections were anteroposterior (AP): −2.4 mm and mediolateral (ML): −1.4 mm relative to the Bregma and dorsoventral (DV): −4.3 mm from the dural surface, calculated according to the mouse atlas of Franklin and Paxinos.^59^ One µl of AAV9-TSS2-sg2 or AAV9-control sgRNA with a titer of 1.50 x10^10^ GC/ µl were injected unilaterally onto the right substantia nigra using a 5 μL Hamilton syringe at a speed of 0.1 µl/15 sec. The needle was left in place for an additional 5 min before it was withdrawn slowly from the brain parenchyma. Skull was cleaned with a sterile gauze and the incision was closed with 1-2 stitches using a 5-0 non-absorbable suture (ETHICON. VCP493G). Buprenorphine SR (0.5 mg/ml) was administered as post-surgical analgesia and wound was sanitized with a 70% ethanol wipe and iodine solution twice. Mice were monitored after surgery and recovered in a warm recovery cage until full motor recovery.

One month and six months after surgery, animals were euthanized with isoflurane (induction 4%, euthanasia 6-8 %) and cardiac perfusion was performed with 100 ml of 4 C PBS (0.1 M, pH 7.2) placing a 25G 0.50 mm X 16 mm syringe (BD Vacutainer, 367294) through the left ventricle of the heart to remove blood from the brain and organs. Brain was harvested and placed onto a rectangular mold (peel away disposable embedding mold rectangular, 22×30 mm), embedded with OCT indicating frontal and posterior part of the brain with a lab marker and fast frozen in liquid nitrogen.

### Preparation of tissue sections for in situ hybridization (RNAscope) and immunostaining

Before tissue sectioning, the frozen brain was taken from −80°C to a cryostat (ThermoScientific, HM525NX) for at least 2 hours to equilibrate to −14°C (sectioning temperature). The tissue block was peeled away from the mold and grossly sectioned with a blade into 3 sections: frontal (olfactory bulb to the beginning of the hippocampal formation), medial (hippocampal formation to beginning of cerebellum), and posterior (cerebellum). The mesencephalon (medial section) between bregma −2.7 to −3.9 was sectioned in 20 µm thick coronal sections and placed onto SuperFrost Plus slides (Fisher Scientific # 12-550-15). Three consecutive sections were placed per slide and transferred to −80°C until further processing.

### RNAscope Multiplex Fluorescent Assay (ACD Bio, Cat. No. 320850)

*In situ* hybridization RNAScope was performed according to manufacturer’s instructions using the RNAscope multiplex fluorescent assay v.2 kits (ACD Systems, Cat. No. 323100). In brief, slides were selected (on dry ice) and taken from storage temperature (−80°C) to pre-chilled 10% neutral buffered formalin for 15 min at 4°C. After fixation, tissue sections were dehydrated gradually in 50% ethanol (5 min, RT), in 70% ethanol (5 min, RT), and then in twice in 100% ethanol (5 min, RT). Sections were air-dried for 5 min after ethanol treatment. Tissue sections were treated with RNAScope hydrogen peroxide (Part Number 322335) for 10 min at RT, followed by a protease IV incubation (10 min, RT) inside a Hybez humidity control tray wet with distilled water. Hybridization probes: Hs-*SNCA* (ACD Systems, Cat. No. 605681-C1) and, sadCas9-C3 (1:50) (ACD Systems, Cat. No. 501621-C3), positive control (ACD Systems, Cat. No. 320861) and negative control (ACD Systems, Cat. No.320871) were performed in a hybridization oven (ACD, HybEZ™ II Hybridization System, ACD Systems, Cat. No. 321710) at 40°C for 2 hours. Slides were washed and underwent three steps of amplification: AMP1 (30 min, 40°C), AMP2 (40°C), and AMP3 (15 min, 40°C). Then, slides were incubated with RNAScope HRP-C1 for 15 min at 40°C followed by a washing cycle and incubation with TSA Plus fluorophore (1:600) for 30 min at 40°C. Lastly, slides were blocked with a horse radish peroxidase (HRP) blocker for 15 min at 40°C and counterstained with a nuclear marker (DAPI). RNAScope slides were covered with Prolong Gold Antifade reagent (Invitrogen, P36930) and a 22x 50 mm coverslip, then sealed with nailpolish before imaging.

Imaging was performed on ImageXpress Pico (Molecular Devices) with a 20X air objective with an image acquisition software (Molecular devices version 2.6.130). For protein acquisition imaging protocol, the exposure used was 10, 300 and 200 ms for 405, 555 and 647 nm fluorophores respectively. A Z-stack acquisition mode with 2 planes and a focus step of 5 um under a “Best focus” acquisition mode was used. For RNA acquisition imaging protocol, the exposure used was 20, 100 and 200 ms for 405, 555, and 647 fluorophores respectively. A Z-stack with 3 planes and focus step of 2 μm was used under a “maximun fluorescence” acquisition mode. After acquisition, stitched images of each individual channel (fluorophore) are exported as “TIFF” files for image analysis.

### Immunofluorescence

Tissue sections were transferred from −80°C immediately to a 10% formalin solution at room temperature, neutral buffered (Sigma-Aldrich, HT501128-4L) for 15 min, washed with PBS and permeabilized with 0.3 % Triton X-100 for 20 minutes. Blocking was performed with 5% goat serum and 1% bovine serum albumin for 30 minutes at room temperature. Overnight incubation with primary antibodies at 4°C was performed using the following antibodies: Cas9 (Diagenode, C15200230-100, 1:500), alpha synuclein (Abcam, MJFR1-ab138501, 1:500), IBA1 (Fujifilm, 019-19741, 1:500), NeunN (SySy, 266008, 1:500). After primary incubation, slides were washed 3 times for 10 minutes each. Alexa fluor 488, 550, or 647 nm were used as fluorophores. Counterstain with Hoechst (LifeTechnologies, H3570) was used to labeled nuclei and slides were mounted into coverslips with ProLong Antifade Mountant media (ThermoFisher Scientific, P36970).

### QuPath image analysis pipeline

#### SNCA mRNA quantification

Stitched individual images of the three different channels (fluorophores); DAPI, Cas9 mRNA and *SNCA* mRNA were merged using ImageJ. Merged pictures are then imported into a QuPath project. Regions of interest (ROI) are selected using a polygonal hand free drawing tool. One ROI was drawn per image which include the substantia nigra pars compacta and ventral tegmental area from the injected side. This region was then analyzed using a two-step pipeline for image analysis; 1. A “Cell detection” algorithm with the following settings: background radius (15px), Median filter radius (2px), Sigma (10 px), minimum area (10 px^2^), Maximum area (1000 px^2^), threshold (5), Cell expansion (40 px). 2. A “subcellular spot detection” was performed using the following parameters: Detection threshold (150), smooth before detection, split by intensity, split by shape with an expected spot size of (30 px^2^), minimum spot size (4 px^2^) and a maximum of (40 px^2^). Results of number of mRNA foci were reported in QuPath and then exported as an Excel file for further treatment and processing. In Excel: Normalization and comparison between Bregma levels was performed using the ROI measurements from the injected side in all the groups as followed:

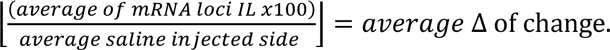

For comparison between groups the relative expression was compared and normalized with saline group percentages and results were plotted into GraphPad for statistical analysis.

#### Alpha-synuclein protein quantification

Stitched individual images of the three different channels (fluorophores); DAPI, Cas9 mRNA and *SNCA* mRNA were merged using ImageJ. Merged pictures are then imported into a QuPath project. Regions of interest (ROI) are selected using a polygonal hand free drawing tool. One ROI was drawn per image surrounding the *Substantia nigra pars compacta* and *ventral tegmental area* from the injected side.

Pipeline for image analysis consists in a single process; 1. A “Cell detection” algorithm with the following settings: background radius (15px), Median filter radius (2px), Sigma (10 px), minimum area (10 px^2^), Maximum area (1000 px^2^), threshold (5), Cell expansion (40 px). This algorithm generated intensity measurements of each individual cell a-syn intensity, cell maximum and minimum intensities. This information is the exported into excel files to filter the necessary information. In Excel, maximum intensity is subtracted to the minimum intensity, this generates a range of intensity per cell. Range results are normalized with the mean value of the saline control group. These results were inserted and plotted into GraphPad for statistical analysis.

#### GraphPad statistics

Delta of change per biological replicates and technical replicates where imported as column comparison. A Kruskal-Wallis test for multiple comparison was used to compare the “Saline” group vs the sgRNA and the TSS2 group. Results from statistical analysis are shown as followed: *ns* = *P* > 0.05,∗ = *P* ≤ 0.05, ∗∗ = *P* ≤ 0.01, ∗∗∗ *P* ≤ 0.001.

## Acknowledgments

The study was funded by the Michael J. Fox Foundation, Therapeutic Pipeline Program (MJFF-17123).

The authors declare no conflict of interest. U.S. Provisional Patent Application No. 63/321,035 was filed by Stanford University. L.S.Q. is a Chan Zuckerberg Biohub – San Francisco investigator.

## Author contributions

C.A.M.T., F.Z. implemented study, performed in vivo experiments, analyzed data, and wrote initial draft sections; D.S., D.P. supported design of vectors and in vitro assessments; D.C.H. assisted with histology and immunostaining; L.S.Q. provided CRISPR plasmid and technical support; D.K. supported data interpretation and manuscript editing; B.S. conceptualized the study, obtained financial support, provided oversight on planning and execution, interpreted data, and wrote the manuscript. All authors read, provided feedback, and approved the final version of the manuscript.

## Supplemental Material

**Supplemental Figure 1.**
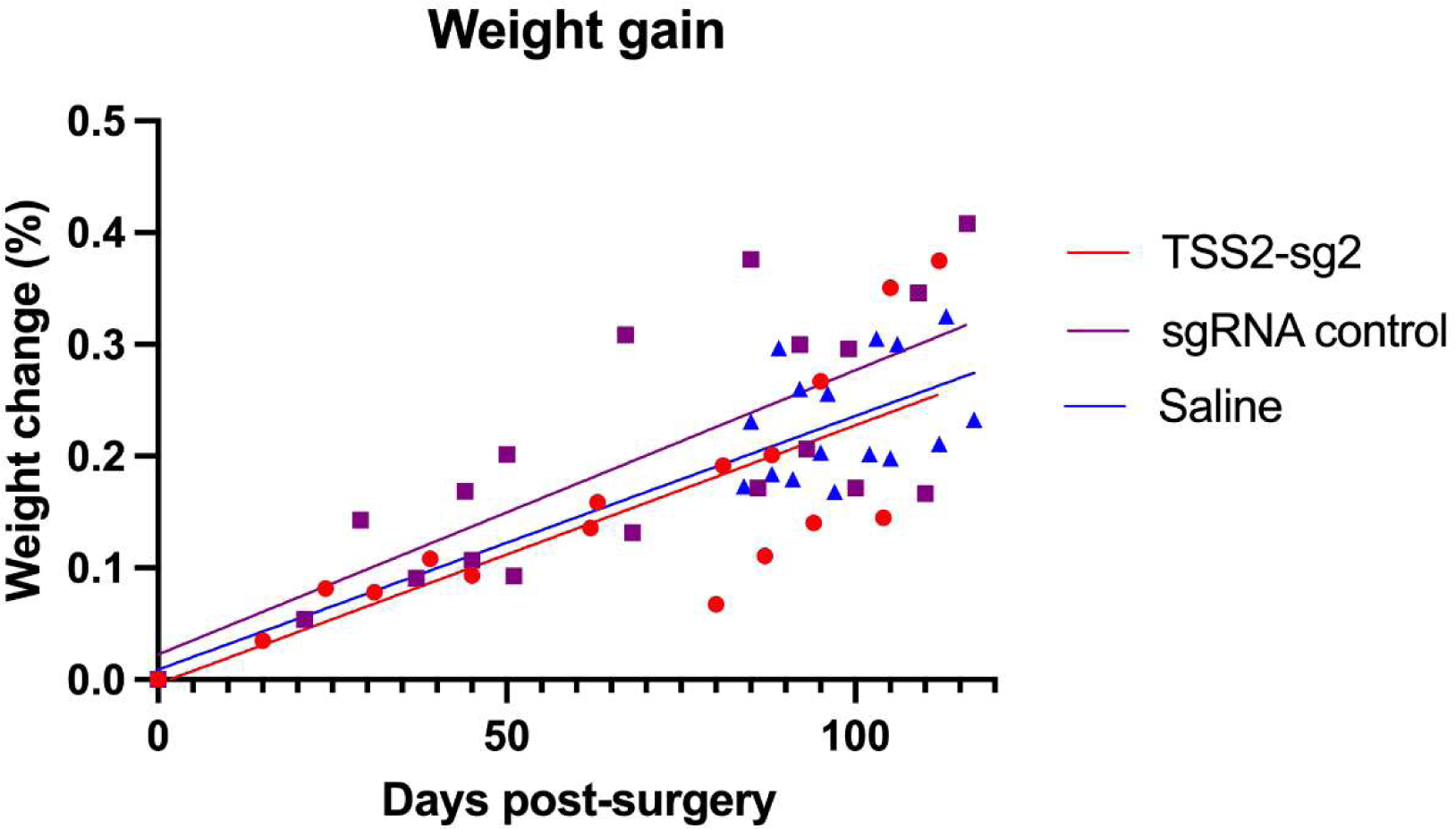
Weight gain of mice post-surgery. All mice gained weight after surgery and we did not detect a statistically significant difference between the three experimental groups.

**Supplemental Figure 2.**
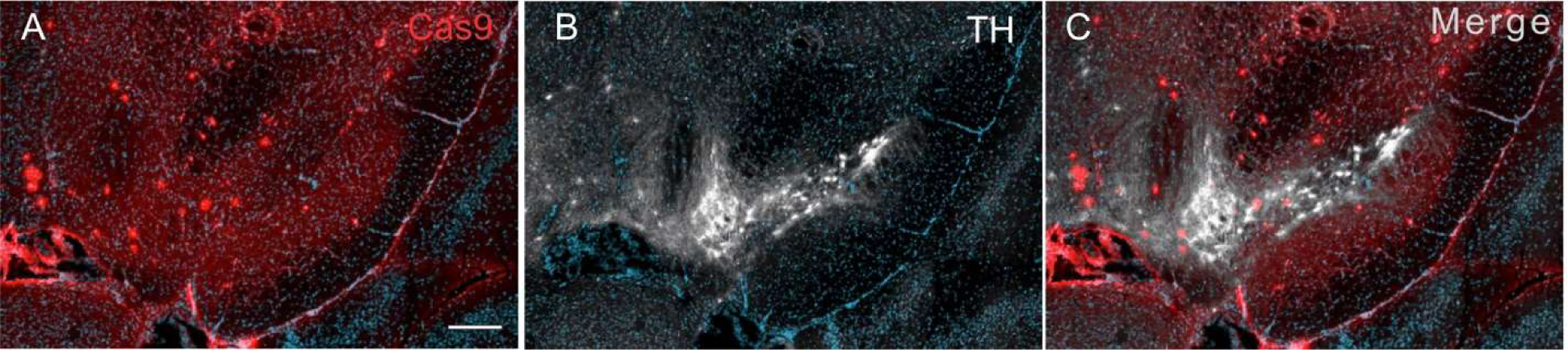
Cas9 and tyrosine hydroxylase (TH) immunostaining in substantia nigra 1 month post stereotactic surgery. **(A)** Cas9 is observed in dopaminergic regions like the ventral tegmental area and substantia nigra pars compacta, **(B)** shows positive immunostaining for TH, **(C)** merged image. Scale bar represents 200 μm.

**Supplemental Figure 3.**
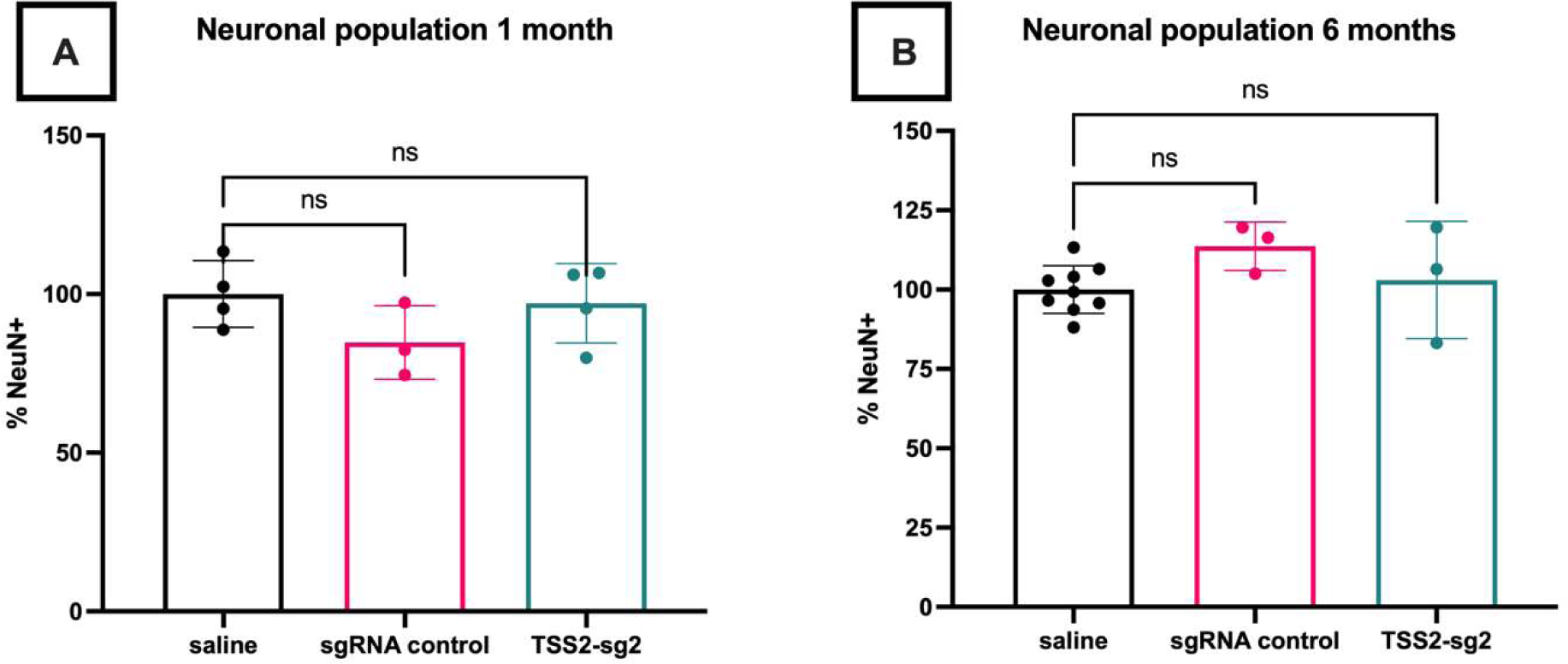
No change on neuronal cell count in the Substantia nigra pars compacta and Ventral Tegmental area at 1- and 6-months post-injection. Graphic representation of neuronal population labeled by immunofluorescence of NeuN in the Substantia nigra and Ventral Tegmental Area at 1 month (**A**) and 6 months (**B**). No statistical difference was observed between groups and times. Saline (n=2), sgRNA control (n=1), TSS2-sg2 (n=1). Results are shown as ns = non-significant.

